# Effects of aging on successful object encoding: Enhanced semantic representations compensate for impaired visual representations

**DOI:** 10.1101/2022.12.10.519871

**Authors:** Loris Naspi, Charlotte Stensholt, Anna E Karlsson, Zachary A Monge, Roberto Cabeza

**Affiliations:** Department of Psychology, Humboldt University of Berlin, Unter den Linden 6, 10117 Berlin, Germany; Center for Cognitive Neuroscience, Duke University, Durham, NC 27708, USA

**Keywords:** episodic memory, aging, fMRI, memory encoding, representational similarity analysis, compensation

## Abstract

Whereas episodic memory and visual processing decline substantially with healthy aging, semantic knowledge is generally spared. There is evidence that older adults can take advantage of their spared semantic knowledge to support their performance in episodic memory and visual tasks. Here, we used fMRI combined with representational similarity analyses (RSA) to examine how visual and semantic representations stored during encoding predict subsequent object memory. Young and older adults encoded images of objects during fMRI scanning and recalled these images while rating the vividness of their memories. After scanning, participants discriminated between studied images and similar lures. RSA based on a deep convolutional neural network and normative concept feature data was used to link patterns of neural activity during encoding to visual and semantic representations. The quality of visual representations was reduced in older adults, consistent with dedifferentiation, whereas the quality of semantic representations was enhanced in older adults, consistent with hyperdifferentiation. Despite dedifferentiation, visual representations stored in early visual cortex predicted later recall with high vividness in both young and older adults, with no age-related differences. In contrast, semantic representations in lingual and fusiform gyrus were associated with better subsequent object picture recall in older but not in young adults. This finding is consistent with evidence that older adults rely on semantic knowledge to compensate for cognitive deficits. Taken together, the results suggest that the age-related neural dedifferentiation for visual information in posterior regions might be partly counteracted by a boost on semantic representations in more anterior areas.

**Significance Statement:** Previous research has shown that healthy aging tends to impair memory for individual events, visual processing, and other cognitive abilities but not semantic knowledge. We investigated the effects of aging on the quality of the information stored in the brain when viewing common objects and on how this information enables subsequent memory for these objects. Using fMRI combined with modeling of the stimuli, we found that visual information was degraded in older adults, but it was sufficient to support subsequent memory. In contrast, semantic information supported subsequent memory only in older adults. This is the first direct neuroscience evidence that older adults take advantage of spared semantic representations to boost their memory for individual events.

## Introduction

In general, healthy older adults show substantial decline in *episodic memory*—memory for context-specific events—but little or no deficits in *semantic memory*—general knowledge of the world (Hoyer and Verhaeghen, 2006). Perhaps for this reason, age-related episodic deficits are reduced, or even eliminated, when to-be-remembered episodic information is meaningful and fits well with prior semantic knowledge (Umanath and Marsh, 2014) In contrast, older adults have considerable difficulty remembering meaningless sensory stimuli (Monge and Madden, 2016). These results suggest that aging differentially impairs the encoding of visual representations but not the encoding of semantic representations. Although behavioral results are generally consistent with this hypothesis (Castel, 2005; Castel et al., 2013; McGillivray and Castel, 2017), the neural mechanisms are still unclear.

It is well-established that visual object processing progresses along the *occipito-temporal cortex* (OTC), from the analysis of simple visual features in early visual cortex to the analysis of objects’ identities and categories in anterior temporal and ventral frontal areas (Mishkin et al., 1983; Murray and Bussey, 1999; Murray and Richmond, 2001). It is also known that these regions store visual and semantic object representations during object encoding and that the quality of these neural representations predict later object memory (Davis et al., 2021; Naspi et al., 2021). Moreover, we know that aging is associated with a reduction in the specificity of object representations in OTC (Carp et al., 2011; Koen et al., 2019; Trelle et al., 2019), and that this deficit—known as *dedifferentiation* (Koen and Rugg, 2019)*—*correlates with poor cognitive performance in older adults, particularly episodic memory (Park et al., 2010; Du et al., 2016). However, our group recently found that although *early visual cortex* (EVC) showed the expected age-related dedifferentiation in visual representations, the *ventral anterior temporal lobe* (vATL) showed an age-related increase in the differentiation of semantic representations (Deng et al., 2021). This effect, which we called *age-related hyperdifferentiation*, was indirectly linked to memory because vATL also showed greater similarity between encoding and retrieval representations (*encoding-retrieval similarity*, ERS) in older adults than young adults. The current study goes beyond Deng et al. (2021) by directly examining whether the quality of visual and semantic object representations stored during encoding differentially predict subsequent memory in young and older adults. Moreover, it also investigates if older adults’ reliance on semantic representations could be linked to compensation.

Young and older adult participants encoded pictures of common objects paired with their name and, during retrieval, they read the names of studied objects, recalling the associated picture, and rating its quality. These subjective ratings were validated with a forced-choice recognition test outside the scanner. RSA was used to measure the quality of visual and semantic object representations stored during encoding, which was then used to predict subsequent object memory. In addition to the two regions of interest (ROIs) investigated by Deng et al. (2021), EVC and vATL, we also examined the lingual gyrus (LG), the fusiform gyrus (FG), and the lateral occipital cortex (LOC), which are regions known to store object representations and predict subsequent object memory (Davis et al., 2021; Naspi et al., 2021).

We investigated two predictions. First, based on ERS results of Deng et al. (2021), we expected that the visual representations in posterior OTC regions would predict subsequent memory more strongly in young than older adults, whereas semantic representations in anterior OTC areas would predict later memory more strongly in older than young adults. Given that Deng et al. (2021) investigated scenes and the current study examines object stimuli, the specific OTC regions that would show these effects were more difficult to predict. Second, assuming semantic representations can play a compensatory role in older adults, we predicted that, in this group, the strength of semantic representations would be not only be positively correlated with later memory, but also negatively correlated with the strength of visual representations (i.e., individual with weaker visual representations compensates with stronger semantic representations).

## Materials and Methods

### Participants

A total of 45 participants took part in the present study, consisting of 23 young adults aged 18-23 years and 22 older adults aged 65-80 years. Participants self-reported to be right-handed, native English speakers, and to have no significant health problems or are taking medications known to affect cognitive function or cerebral blood flow. The Beck Depression Inventory (BDI; Beck et al., 1961) was used to screen for depression, with an exclusion criterion of > 10. The older adults were screened for dementia using the Montreal Cognitive Assessment (MoCA; Nasreddine et al., 2005), with an exclusion criterion of < 26. Following the exclusion of participants based on the screening measures, technical errors, imaging artifacts, and lack of behavioral responses, a total of 18 young adults (M = 20.94, SD = 3.24; 8 females, 10 males) and 17 older adults (M = 70.5, SD = 3.38; 8 females, 9 males) were included in the present study. The study was conducted and data collected by researchers at Duke University, and all experimental procedures were approved by the Institutional Review Board at Duke University. Informed consent was obtained from all participants prior to testing, and they were given monetary compensation for their time following study completion.

### Stimuli

The stimuli used in the present study consisted of 240 of 995 basic-level concepts used for an online object norming experiment (Hovhannisyan et al., 2021). These were members of 8 superordinate categories (*Birds, Clothing, Fruits, Mammals, Outdoor items, Vehicles, Vegetables, Tools*), and 120 were living and 120 non-living things. Two images for each basic-level concept were identified using image search engines Google Images, Bing Images, and Flickr. One of the two images was encoded and the other was used as a distractor in a two-alternative forced-choice recognition test. The allocation of the two images to each condition (i.e., studied item or similar lure) in the two-alternative forced-choice recognition test was counterbalanced across participants. For a manipulation whose results are not reported here, half of the encoded objects were presented in their original resolution, and half, slightly blurred. All encoded objects were subsequently tested in a free recall task, using the object name as a cue, and then in the two-alternative forced-choice recognition test. Each study and test list were presented in a unique, random trial order.

### Procedure

Following the screening procedure (see Participants section and Table 1), the experiment consisted of two scanning sessions approximately 24 hours later, one for encoding and one for retrieval. Stimuli were presented using E-Prime 3.0 software (Psychology Software Tools, Pittsburgh, PA). In the scanner, stimuli were viewed on a back-projection screen via a mirror attached to the head coil. Earplugs were used to reduce scanner noise, and head motion was minimized using foam pads. At encoding, participants were shown the 240 object images with their accompanying object label. The object was centered on a white background with the object label below. The scanned encoding phase consisted of eight runs, with 30 trials in each run. Each object image and label were presented once. Participants made either a clear-blurry judgment or a living-nonliving judgment, but the results of these conditions were collapsed for the current study to increase statistical power. Each object was presented on the screen for 4 sec, and participants made responses via a keypad. Each encoding trial was followed by an 8 sec active baseline interval, where participants were presented with a number from one to four and asked to press the corresponding keypad button. At retrieval, the runs were identical in order and duration to those at encoding, except that only the object labels (e.g., *table*) from encoding were shown. In each trial, participants recalled the image of the corresponding encoded object in as much detail as possible and rated the vividness (1 = *least amount of detail*, 4 = *highly detailed memory*). After exiting the scanner, participants performed the two-alternative forced-choice task. In each trial, participants saw a studied picture and a similar example of the same object (e.g., a different table), and they had to choose the encoded image and rate their confidence (1 = *low confidence*, 4 = *high confidence*). This task was self-paced.

**Table 1.**
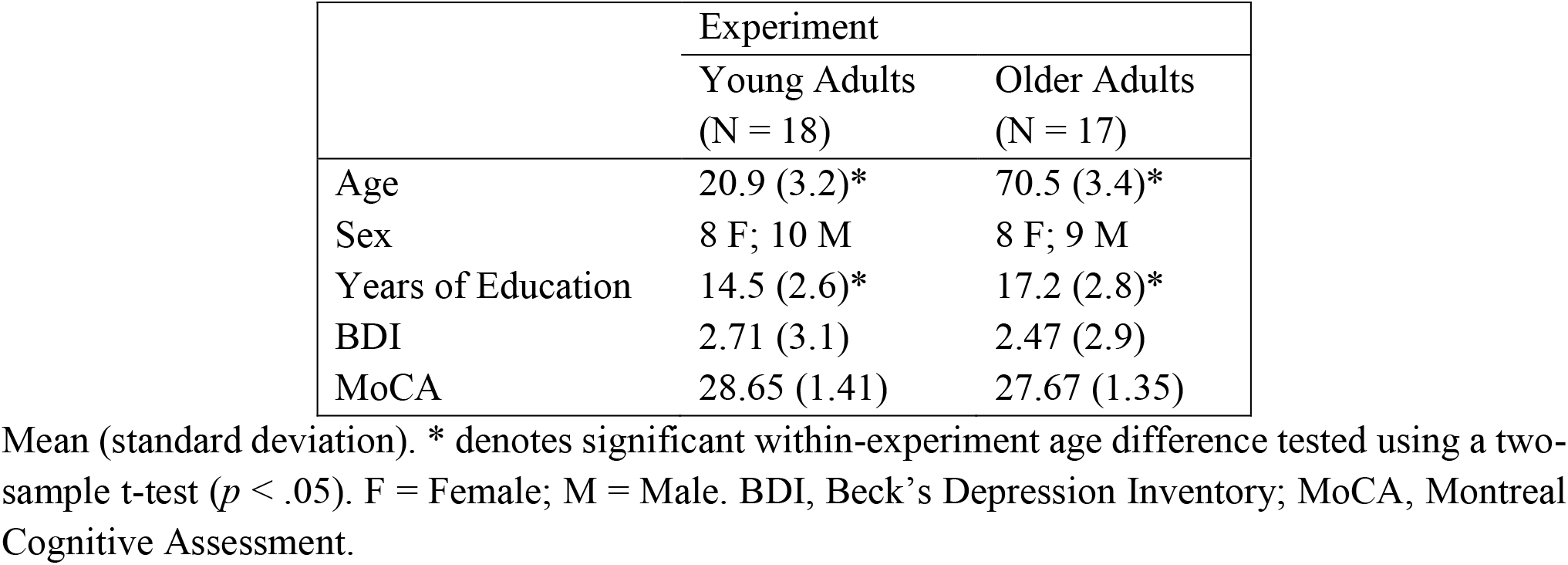
Demographic and Cognitive Tests Data.

|  | Experiment |  |
| --- | --- | --- |
|  | Young Adults<br>(N = 18) | Older Adults<br>(N = 17) |
| Age | 20.9 (3.2)* | 70.5 (3.4)* |
| Sex | 8 F; 10 M | 8 F; 9 M |
| Years of Education | 14.5 (2.6)* | 17.2 (2.8)* |
| BDI | 2.71 (3.1) | 2.47 (2.9) |
| MoCA | 28.65 (1.41) | 27.67 (1.35) |

Mean (standard deviation). \* denotes significant within-experiment age difference tested using a two-sample t-test ( $p < .05$ ). F = Female; M = Male. BDI, Beck's Depression Inventory; MoCA, Montreal Cognitive Assessment.

### fMRI acquisition

The data was collected using a General Electric 3T Premier UHP MRI scanner and a 48-channel head coil. The scanning session started with a localizer scan, followed by a high-resolution anatomical image (162 axial slices parallel to the AC-PC plane, voxel dimension of 1.0 x 1.0 x 1.0 mm^3^). This was followed by a resting-state scan. Whole-brain functional scans were collected using a multiband echo-planar imaging scan (repetition time = 2 sec, echo time = 30 msec, field of view = 256 mm^2^, oblique slices with voxel dimension of 2.0 x 2.0 x 2.0 mm^3^). The stimuli were projected onto a mirror at the back of the scanner bore, and a four-button fiber-optic response box was used to record the participants’ in-scan responses. To minimize scanner noise, participants wore earplugs, foam pads were used to minimize head motion, and MRI-compatible lenses were used to correct vision if needed. At the end of the scan diffusion-weighted images were collected, but together with the resting-state scan they will not be reported on in the present study.

### Image preprocessing

Data processing was performed using *fMRIPrep* (20.2.3) (Esteban et al., 2019). First, each T1-weighted volume was corrected for intensity non-uniformity (INU) with N4BiasFieldCorrection (Tustison et al., 2010), distributed with ANTs 2.3.3 (Avants et al., 2008). The T1w-reference was then skull-stripped with a *Nipype* implementation of the antsBrainExtraction.sh workflow (from ANTs), using OASIS30ANTs as a target template. Brain tissue segmentation of cerebrospinal fluid (CSF), white-matter (WM) and gray-matter (GM) was performed on the brain-extracted T1w using fast (FSL 5.0.9; Zhang et al., 2001). A T1w-reference map was computed after registration of 2 T1w images (after INU-correction) using mri_robust_template (FreeSurfer 6.0.1; Reuter et al., 2010). Volume-based spatial normalization to one standard space (MNI152NLin2009cAsym) was performed through nonlinear registration with antsRegistration (ANTs 2.3.3), using brain-extracted versions of both T1w reference and the T1w template. For each of the 8 BOLD runs found per subject, the following preprocessing was performed. First, a reference volume and its skull-stripped version were generated using a custom methodology of *fMRIPrep*. A B0-nonuniformity map (or fieldmap) was estimated based on two (or more) echo-planar imaging (EPI) references with opposing phase-encoding directions with 3dQwarp (Cox and Hyde, 1997). Based on the estimated susceptibility distortion, a corrected EPI (echo-planar imaging) reference was calculated for a more accurate co-registration with the anatomical reference. The BOLD reference was then co-registered to the T1w reference using flirt (FSL 5.0.9; Jenkinson and Smith, 2001) with the boundary-based registration (Greve and Fischl, 2009) cost-function. Co-registration was configured with nine degrees of freedom to account for distortions remaining in the BOLD reference. Head-motion parameters with respect to the BOLD reference (transformation matrices, and six corresponding rotation and translation parameters) are estimated before any spatiotemporal filtering using mcflirt (FSL 5.0.9; Jenkinson et al., 2002). BOLD runs were slice-time corrected using 3dTshift from AFNI 20160207 (Cox and Hyde, 1997). The BOLD time-series (including slice-timing correction when applied) were resampled onto their original, native space by applying a single, composite transform to correct for head-motion and susceptibility distortions. These resampled BOLD time-series will be referred to as unsmoothed preprocessed BOLD in original space. For RSA, we used these BOLD time-series to keep the finer-grained structure of activity.

### Experimental design and statistical analysis

#### Behavioral analysis

To assess differences in the study phase reaction time as a function of subsequent recall, we used a linear mixed effect model with vividness ratings (subsequent recall memory coded as 3 or 4 and subsequent forgotten memory coded as 1 or 2) and age group (young adults and older adults) as predictors, and reaction time (RT) as the outcome variable. At test, to confirm the validity of in-scan memory vividness ratings as a memory measure, we used a generalized linear mixed effect model to investigate age-related differences in post-scan memory accuracy (0, 1) predicted by the vividness rating during retrieval.

### Multivariate fMRI analysis

#### Overview

The goal of our study was twofold. In the first analysis, we aimed to replicate and extend to older adults the key findings regarding visual and semantic processing irrespective of memory encoding (Clarke and Tyler, 2014; Devereux et al., 2018; Martin et al., 2018). In a second analysis, we investigated whether visual and semantic representations differentially contributed to later recall memory in young and older adults. Although recent studies suggest that these dimensions are a key factor in the young adults (Davis et al., 2021; Naspi et al., 2021), little is known about their contribution to memory in older adults (but see Deng et al., 2021). Our analytical method involved 4 steps. The first 3 steps are standard in RSA studies (Kriegeskorte et al., 2008), and were used for the overall analysis irrespective of memory encoding. 1) The visual and semantic properties of the stimuli were used to create 2 different representational dissimilarity matrices (RDMs). In an RDM, the rows and columns correspond to the stimuli (240 in the current study) and the cells contain values of the predicted dissimilarity (1—Pearson correlation) between pairs of stimuli. 2) A pattern of activity was extracted for each region of interest (ROI). This matrix has the same structure as the RDM (stimuli in rows and columns), but the cells do not contain a measure of dissimilarity in stimulus properties as in the RDM, but dissimilarity in the fMRI activation patterns by the stimuli. 3) After vectorizing both the model RDMs and brain RDMs, we tested the model-brain fit for the object processing analysis irrespective of memory encoding using a one-sample Wilcoxon signed rank test and a two-sample Wilcoxon rank sum test for age-related differences.

For the second analysis, after step 2) we followed a slightly different procedure (see also Davis et al., 2021). 4) We computed the correlation between the dissimilarity of each object with the rest of the objects in terms of stimulus properties (each row of the model RDM) and the dissimilarity of the same object with the rest of the objects in terms of activation patterns (each row of the brain RDM) and identified brain regions that demonstrated a significant correlation across all items and subjects. Davis et al. (2021) termed the strength of this second-order correlation as the item-wise RDM-activity fit (IRAF). The IRAF in a brain region is therefore an index of the sensitivity of that region to that particular kind of visual or semantic representation. Note that such an item-wise approach differs from the typical method of assessing such second-order correlations between brain and model RDMs (Kriegeskorte and Kievit, 2013), which typically relates the entire item × item matrix at once. This item-wise approach is important for linking visual and semantic representations to subsequent memory for specific objects. To test whether these representations predicted successful recall, we ran a generalized linear mixed effect model and post-hoc differences of estimated marginal means. The implementation of the steps for these two analyses is outlined in the following sections.

#### RSA first-level GLM

To perform the representational similarity analyses a beta estimate image for each trial was created. This was conducted in SPM12 using the first-level general linear model (GLM) and a Least-Squares-All (LSA) methods (Mumford et al., 2012). A design matrix was created for each participant which included one regressor for each trial, computed by convolving the 4 sec duration stimulus function with a canonical hemodynamic response function. For each run, six motion regressors were included consisting of the three translations and three rotations estimated during spatial realignment, as well as session constant for each of the 8 runs. The model was fit to native space preprocessed functional images using Variational Bayes estimation with an AR(3) autocorrelation model (Penny et al., 2005). A high-pass filter cut-off of 128 sec was applied, and the data was scaled to a grand mean of 100 across all voxels and scans within sessions. A whole brain-mask was created by including voxels which had at least a 0.1 probability of being in grey or white matter, as indicated by the tissue segmentation of the participants’ T1 scan.

#### ROIs

The ROIs used in the current study are shown in Figure 1. These regions were selected because they significantly predicted subsequent memory in two recent studies with young adults (Davis et al., 2021; Naspi et al., 2021) and age-related differences in memory reactivation (Deng et al., 2021). We defined five ROIs, including areas spanning the posterior and anterior ventral stream, which have been implicated in visual and semantic feature-based object recognition processes (Clarke and Tyler, 2014, 2015; Lambon Ralph et al., 2017; Devereux et al., 2018). Except where explicitly stated, ROIs were bilateral and defined in MNI space using the Harvard-Oxford structural atlas: (1) the early visual cortex (EVC; BA17/18) ROI was defined using the Julich probabilistic cytoarchitectonic maps (Amunts et al., 2000) from the SPM Anatomy toolbox (Eickhoff et al., 2005); (2) the lingual gyrus (LG); (3) the fusiform gyrus (FG); (4) the lateral occipital cortex (LOC); (5) the ventral anterior temporal lobe (vATL) ROI included voxels with .30% probability of being in the anterior division of the inferior temporal gyrus (ITG) and .30% probability of being in the anterior division of the FG. The ROIs in Figure 1 are mapped on a pial representation of cortex using the Connectome Workbench (https://www.humanconnectome.org/software/connectome-workbench).

**Figure 1.**
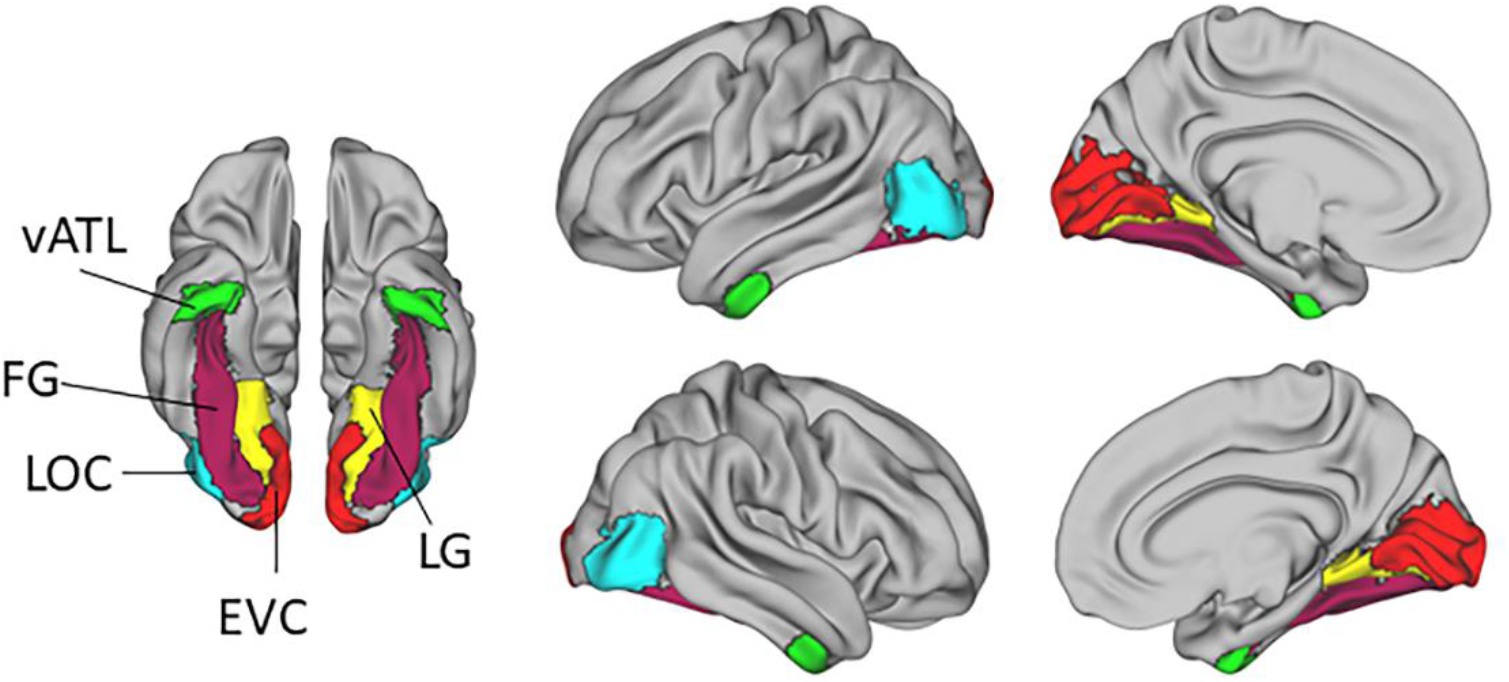
Binary ROIs overlaid on a pial cortical surface based on the normalized structural image averaged over participants. Colored ROIs represent regions known to be engaged in episodic encoding and in visual or semantic cognition (for details, see ROIs).

### RSA ROI analysis

#### Model RDMs

We created two theoretical RDMs using low-level visual and object-specific semantic feature measures. Figure 2 illustrates the multidimensional scale plots for the visual and semantic relations expressed by these models, the corresponding model RDMs, and the correlation between the RDMs.

1. The visual model was derived using a pretrained version of VGG16 from the visual geometry group (Simonyan and Zisserman, 2015), a widely used deep convolutional neural network (DNN). The VGG16 consists of 16 layers, thirteen of which are convolutional layers and three of which are fully connected layers. Evidence suggests that DNNs can simulate the processing that occurs in the ventral visual pathway, where earlier layers detect lower-level visual features and later layers detect higher-level visual features (Zeiler and Fergus, 2014). In the present study we extracted features from the second convolutional layer (layer 2), as a previous study found this layer to approximate low-level visual features such as orientation and edges better than layer 1 (Bone et al., 2020). To extract activation values from layer 2 we used a Python toolbox (Muttenthaler and Hebart, 2021). The visual model was created by feeding the object images through the DNN, extracting the activation values from layer 2, and correlating the values between the stimuli resulting in a 240 x 240 similarity matrix. This was then turned into a dissimilarity matrix by computing the pairwise dissimilarity values as 1 – Pearson’s correlation, resulting in the final visual model.
2. Construction of the semantic feature RDM followed Clarke and Tyler (2014) but used updated property norms (Hovhannisyan et al., 2021). We first computed pairwise feature similarity between concepts from a semantic feature matrix in which each concept is represented by a binary vector indicating whether a given feature is associated with the concept or not. Pairwise dissimilarity between concepts was computed as 1 – S where S is equal to the cosine angle between feature vectors. This RDM captures both categorical similarity between objects (as objects from similar categories have similar features) and within-category object individuation (as objects are composed of a unique set of features). We excluded taxonomic features as they reflect category-level, but not concept-specific, information (Taylor et al., 2012). Analyses were implemented using custom MATLAB version 2021a (The MathWorks) and Python version 3.8.

**Figure 2.**
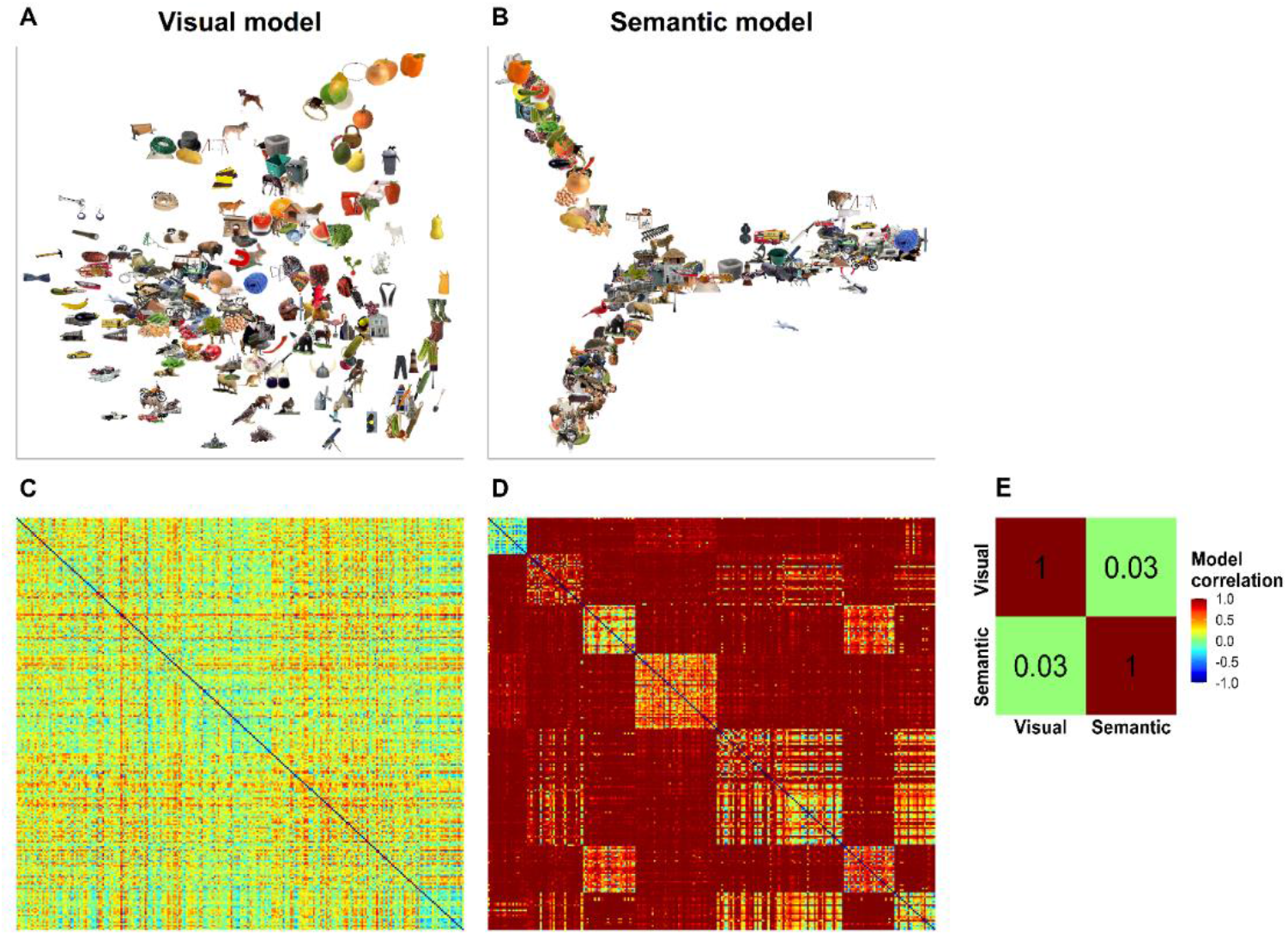
Multidimensional scale plots, model RDMs for visual and semantic similarities, and model correlations. Pairwise dissimilarities were calculated to create RDMs. A, Visual dissimilarity codes for a combination of orientation and edges (e.g., round objects toward the top-right, horizontal shapes on the left, vertical shapes on the bottom-right). B, Semantic dissimilarity codes for finer-grained distinctions based on features of each concept (e.g., fruit and vegetables on top-left, nonliving things on the middle-right, and many categories of animal on the bottom-left). C, Visual dissimilarity model of object processing including all the items. D, Semantic dissimilarity model of object processing including all the items. E, Spearman’s correlation of visual and semantic RDMs.

#### Brain RDMs

In addition to the model RDMs describing feature dissimilarity, we also created brain RDMs, or activity pattern matrices, which represent the dissimilarity in the voxel activation pattern across all stimuli. Thus, the activation pattern matrices have a dissimilarity structure as the RDM with stimuli as rows and columns. However, whereas each cell of an RDM contains a measure of dissimilarity in stimulus’ properties, each cell of an activity pattern dissimilarity matrix contains a measure of dissimilarity in activation patterns across stimuli. As noted above, the activation patterns were extracted for 5 ROIs and correlated with Pearson’s r. As with the model RDMs, each matrix was then transformed into brain RDMs by calculating 1 – Pearson’s correlation.

#### Fitting model RDMs to brain RDMs

For the overall analysis, for each subject and for each ROI, the brain RDM was compared with the model RDM using Spearman’s rank correlation, and the resulting dissimilarity values were Fisher-transformed. Then, for each group separately, we tested for significant positive similarities between model RDMs and brain RDMs using a one-sample Wilcoxon signed rank test. We then tested age-related differences in visual and semantic processing using a two-sample Wilcoxon rank sum test. These tests provide valid inference and treat the variation across subjects as a random effect, thus supporting inference to the population (Nili et al., 2014). We applied a false discovery rate correction for the 5 ROIs. For the subsequent memory analysis, each model RDM was correlated with the activation pattern dissimilarity matrix of each item, in each ROI to obtain an IRAF measure for each item, in each region. Spearman’s rank correlations values were Fisher transformed and mapped back to each region-of-interest. We then used the IRAF as an independent variable in a generalized linear mixed effect model to predict subsequent vividness. The IRAF from the visual and semantic RDMs was used as a continuous predictor in a single model. Thus, the model comprised the IRAF from the visual and semantic model in each of the 5 ROIs for both young and older adults (IRAF × ROIs × group× model). The binary outcome variable was memory indexed as 0 (least number of details recalled, rated as 1 or 2 for vividness) or 1 (maximum number of details recalled, rated as 3 or 4 for vividness). Thus, we measured the predictive effect of each model term by examining the t-statistics for the fixed effect based on beta estimates for each of the 5 ROIs (EVC, LG, FG, LOC, vATL) in addition to the groups of interest (young adults, older adults) and the type of model (visual model, semantic model). Subjects and stimuli were both also entered as random effects. Since the results of these analyses are constrained by the chosen reference-level, we ran post-hoc tests to assess the interaction of our continuous predictor (i.e., IRAF) with each factor (i.e., ROIs, group, and model) on memory by calculating the estimated marginal means. Thus, we derived an estimation for each group in each ROI, and a pairwise comparison of estimated marginal means between groups. These analyses were performed using the lme4 and emmeans packages in RStudio version 1.3.1093.

#### Testing the compensatory role of regions supporting semantic cognition in older adults

After the analysis of the visual and semantic representations involved in object processing and subsequent memory, we tested our hypothesis that regions observed to support semantic cognition may play a compensatory role by boosting older adults’ memory. As noted in the Introduction, this hypothesis is supported by a finding that semantic representations have a stronger effect on subsequent memory in older than young adults. Additionally, this hypothesis predicts a negative correlation between the strength of semantic representations and the strength of visual representations (i.e., individual with weaker visual representations compensates with stronger semantic representation). To test this hypothesis, for each group separately, we performed across-subjects Pearson’s correlation between the model-brain fit (i.e., Spearman correlation coefficient) of regions that revealed *age-related dedifferentiation* of visual representations with the model-brain fit (i.e., Spearman correlation coefficient) of regions that revealed *age-related hyperdifferentiation* of semantic representations.

## Results

### Memory task performance

The linear mixed model at encoding did not reveal any difference in RT related to subsequent old items that were later remembered with high relative to low vividness in the young adults (β = 26.17, SEM = 16.43, *t* = 1.592, *p* = 0.111). Similarly, there was no difference in RT related to subsequent old items that were later remembered with high relative to low vividness in the older adults (β = −9.35, SEM = 17.50, *t* = −0.534, *p* = 0.593). However, older adults showed slower RT compared to young adults for those items that were later recalled with both low vividness (β = 322.91, SEM= 103.62, *t* = 3.116, *p* = 0.002), and high vividness (β = 287.00, SEM = 104.00, *t* = 2.765, *p* = 0.006). We then investigated the memory performance of the two age groups (see Figure 4). At retrieval, confirming the validity of in-scan memory vividness ratings as a memory measure, the generalized linear mixed effect model showed that post-scan memory accuracy was positively related to vividness rating, as is indicated by a significant effect of vividness in both young (β = 0.150, SEM = 0.04, z = 3.808,*p* < 0.001) and older adults (β = 0.206, SEM = 0.04, *z* = 4.648, *p* < 0.001). However, there was no age-related difference in the post-scan memory accuracy as a function of vividness ratings (β = 0.057, SEM = 0.06, *z* = 0.978, *p* = 0.328).

**Figure 3.**
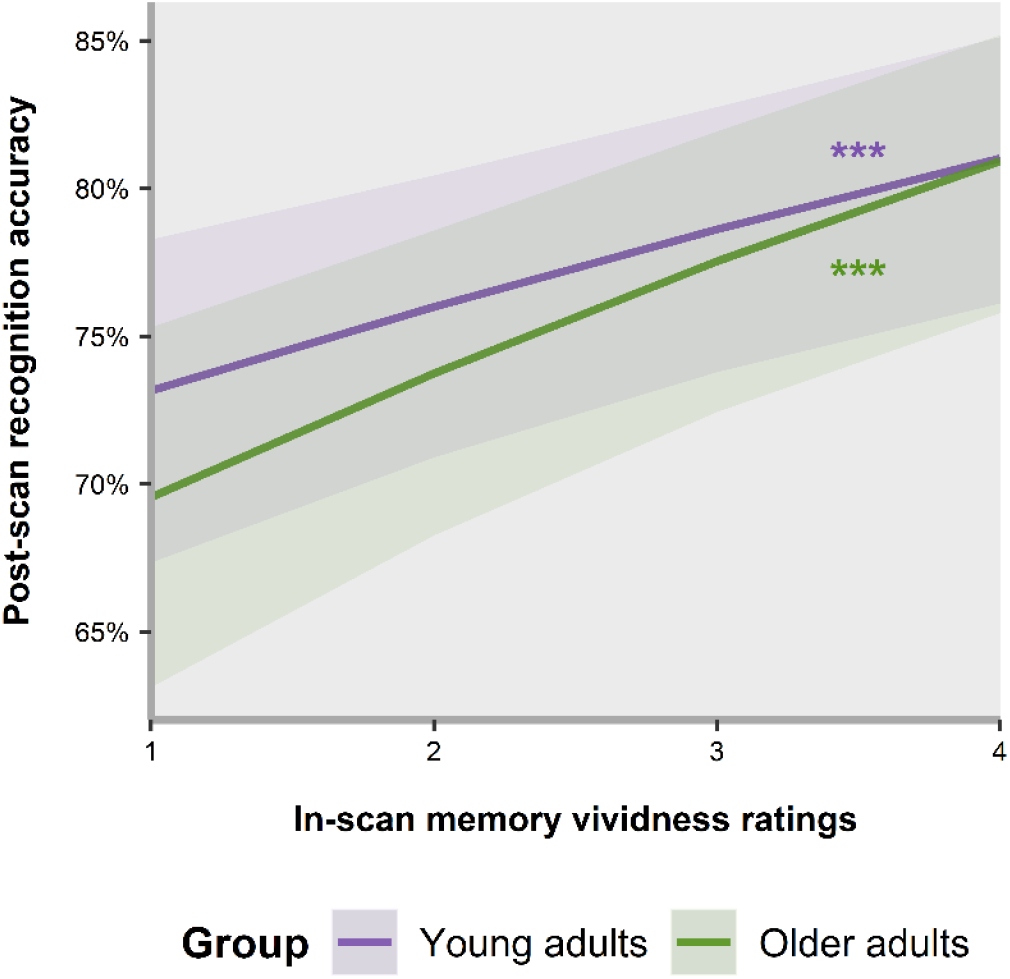
The line plot shows for each in-scan vividness rating value, the corresponding post-scan memory accuracy (hit rate). Shaded colours around the line plots indicate confidence levels (CI) across subjects. Purple and green asterisks indicate *p* values for significant estimated marginal means of linear trends for each group. \**p* < 0.05. \*\**p* < 0.01. \*\*\**p* < 0.001.

**Figure 4.**
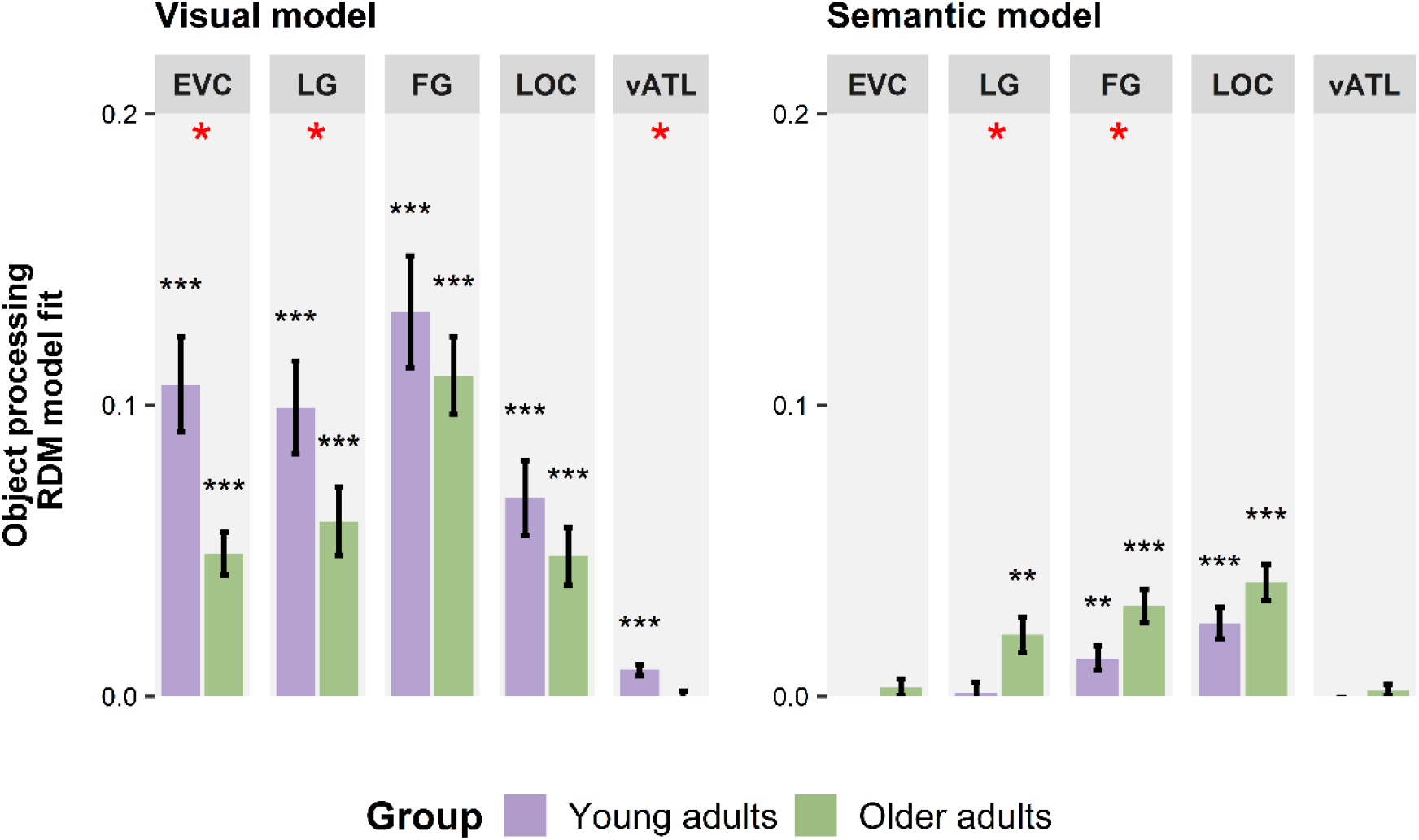
Visual and semantic representations represented in ROIs regardless of memory encoding. Plots represent the strength of visual and semantic representations at the group level within patterns of activity along the ventral stream. Error bars indicate SEM across subjects. Black asterisks above the bars indicate *p* values for tests of whether each individual Spearman’s correlation is > 0 (one-sided Wilcoxon signed rank test; false-discovery rate (FDR) correction calculated across the number of ROIs, i.e., 5). Red asterisks indicate *p* values for age-related differences in Spearman’s correlation (two-sided Wilcoxon rank sum test; false-discovery rate (FDR) correction calculated across the number of ROIs, i.e., 5 per group). \**p* < 0.05. \*\**p* < 0.01. \*\*\**p* < 0.001.

### Visual and semantic object processing irrespective of memory encoding

We first identified brain regions that coded for visual and semantic object representations during perception, independent of memory encoding. As in the studies by Clarke and Tyler (2014) and Naspi et al. (2021), this was done by including all trials in RSA analyses of model-brain regardless of their subsequent memory status. As illustrated by Figure 5, a one-sample Wilcoxon signed-rank test revealed that in young adults, visual representations were reliably represented in all 5 ventral pathway ROIs, including EVC (T = 171, *p* < 0.001), LG (T = 169, *p* < 0.001), FG (T = 169, *p* < 0.001), LOC (T = 165, *p* < 0.001), and vATL (T = 162, *p* < 0.001). Older adults displayed significant visual representations in EVC (T = 153, *p* < 0.001), LG (T = 152, *p* < 0.001), FG (T = 153, *p* < 0.001), and LOC (T = 142,*p* < 0.001) but not in vATL (T = 66, *p* = 0.694). As predicted, the two-sample Wilcoxon sum rank test also revealed age-related differences in the quality of visual representations. More specifically, older adults showed decreased quality of visual representations in EVC (T = 69, *p* = 0.010), LG (T = 95, *p* = 0.048) and vATL (T = 79, *p* = 0.018), whereas visual representations in FG and LOC (respectively, T = 118, *p* = 0.161 and T = 128, *p* = 0.212) were of similar quality as in young adults.

**Figure 5.**
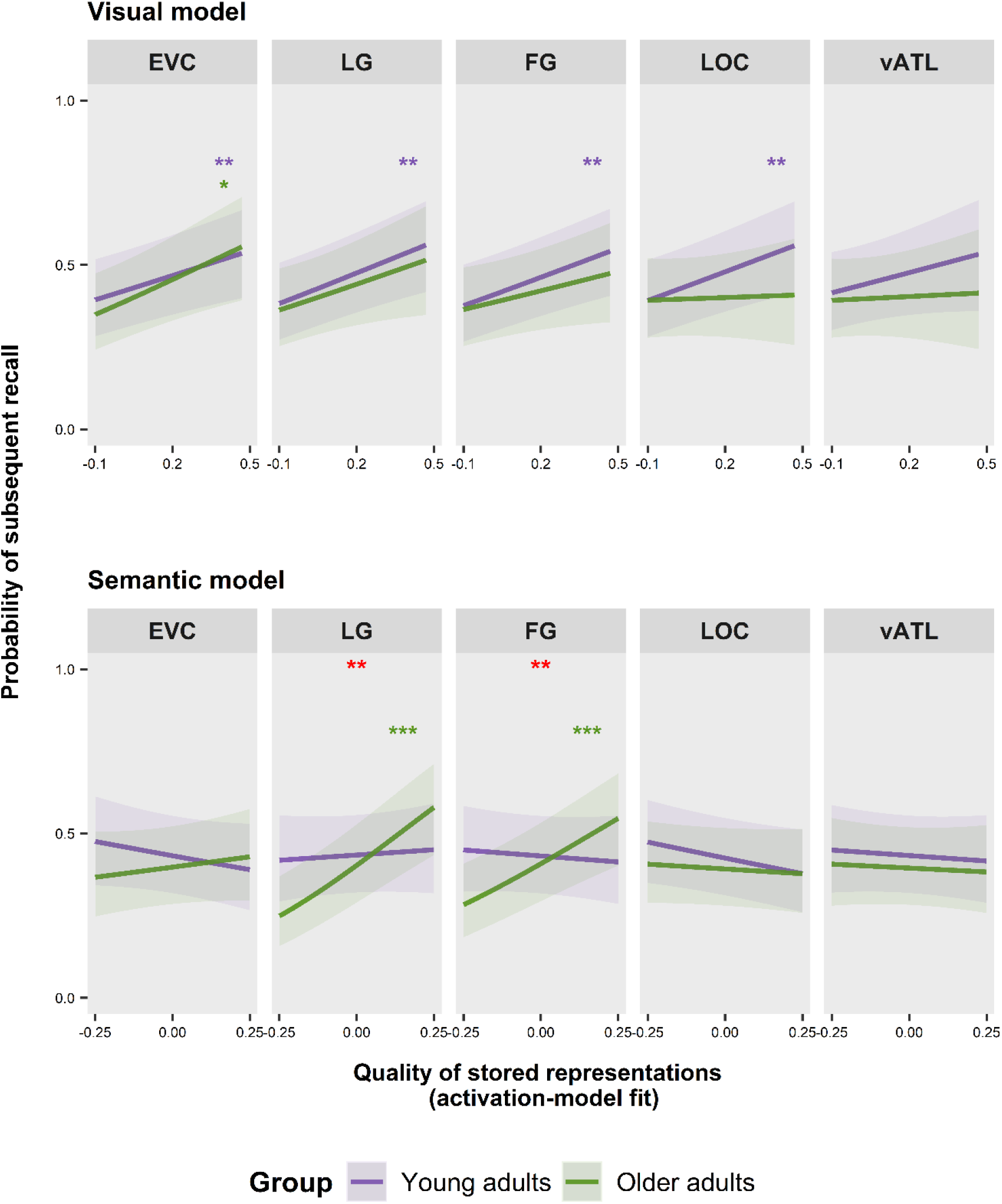
The line plots represent the effect of visual and semantic representations on the probability of recalling studied items with high vividness on young (purple lines) and older (green lines) adults. Shaded colours around the line plots indicate SEM across subjects. Purple and green asterisks indicate *p* values for significant estimated marginal means of linear trends for each ROI in each group; false-discovery rate (FDR) correction calculated across the number of ROIs, i.e., 5 per group. Red asterisks within the middle top of the panels indicate significant age-related differences (difference between estimated marginal means of linear trends; false-discovery rate (FDR) correction calculated across the number of ROIs, i.e., 5 per contrast between groups). \**p* < 0.05. \*\**p* < 0.01. \*\*\**p* < 0.001.

Turning to semantic representations, the one-sample Wilcoxon signed-rank tests revealed that in young adults, semantic object features were significantly coded in LOC (T = 164, *p* < 0.001) and, more anteriorly, in FG (T = 147, *p* = 0.008). None of the other three ROIs showed significant effects in young adults (respectively, T = 45, *p* = 0.963 for EVC; T = 95, *p* = 0.585 for LG; T = 57, *p* = 0.963 for vATL). Like young adults, older adults showed reliable semantic representations in LOC (T = 153, *p* < 0.001) and FG (T = 148, *p* < 0.001), but they additionally displayed significant semantic representations in LG (T = 136, *p* = 0.003). As in young adults, EVC (T = 88, *p* = 0.306), and vATL (T = 92, *p* = 0.305) did not show significant semantic representations in older adults. As for visual representations, a two-sample Wilcoxon sum rank test was used to compare the quality of semantic representations in the two groups. In sharp contrast with the results of visual representations, older adults showed stronger semantic representations than young adults. This effect was found in LG and FG (respectively, T = 227, *p* = 0.018 and T = 231, *p* = 0.018), but not in the other 3 ROIs (respectively, T = 206, *p =* 0.060 for EVC; T = 204, *p* = 0.006 for LOC; T = 200, *p* = 0.063 for vATL).

In each ROI, we also checked which representations showed unique effects after controlling for effects of the other model using Spearman’s partial correlation. However, given the low correlation between models (see Figure 2E), the results of the analysis revealed an identical pattern of findings and, thus, are not reported further. In sum, visual representations were of higher quality in young than older adults (in EVC, LG, and vATL), consistent with the idea of *dedifferentiation* (Koen and Rugg, 2019), whereas semantic representations showed the opposite effect and were stronger in older adults than young adults (in LG and FG), consistent with the idea of *hyperdifferentiation* (Deng et al., 2021). This conclusion is supported by a two-way ANOVA that revealed a significant interaction between group (young adults, older adults) and type of model (visual, semantic) (F(1,346) = 17.5, *p* < 0.001).

### Effect of stored visual and semantic representation on subsequent memory

To examine whether visual and semantic representations at encoding predicted later recall of the object pictures, we ran a generalized linear mixed effect model. The results of the estimated marginal means derived from the full model are reported below in Table 2 and displayed in Figure 5 (upper panel and bottom panel).

**Table 2.** Estimated marginal means for the visual and semantic models separated by age group.

| Visual model |  |  |  |  |
| --- | --- | --- | --- | --- |
| ROI | $\beta$ | SE | z-value | p |
| EVC |  |  |  |  |
| Young adults | 1.01 | 0.32 | 3.11 | 0.002 |
| Older adults | 1.48 | 0.50 | 2.97 | 0.015 |
| LG |  |  |  |  |
| Young adults | 1.27 | 0.39 | 3.28 | 0.002 |
| Older adults | 1.09 | 0.54 | 2.01 | 0.111 |
| FG |  |  |  |  |
| Young adults | 1.18 | 0.34 | 3.43 | 0.002 |
| Older adults | 0.80 | 0.47 | 1.69 | 0.153 |
| LOC |  |  |  |  |
| Young adults | 1.19 | 0.38 | 3.14 | 0.002 |
| Older adults | 0.12 | 0.53 | 0.22 | 0.829 |
| vATL |  |  |  |  |
| Young adults | 0.83 | 0.55 | 1.52 | 0.129 |
| Older adults | 0.16 | 0.64 | 0.25 | 0.829 |
| Semantic model |  |  |  |  |
| ROI | $\beta$ | SE | z-value | p |
| EVC |  |  |  |  |
| Young adults | -0.70 | 0.56 | -1.24 | 0.534 |
| Older adults | 0.52 | 0.57 | 0.91 | 0.607 |
| LG |  |  |  |  |
| Young adults | 0.26 | 0.55 | 0.47 | 0.636 |
| Older adults | 2.85 | 0.58 | 4.93 | <0.001** |
| FG |  |  |  |  |
| Young adults | -0.30 | 0.53 | -0.57 | 0.636 |
| Older adults | 2.23 | 0.54 | 4.09 | <0.001** |
| LOC |  |  |  |  |
| Young adults | -0.79 | 0.42 | -1.86 | 0.312 |
| Older adults | -0.24 | 0.39 | -0.62 | 0.669 |
| vATL |  |  |  |  |
| Young adults | -0.27 | 0.55 | -0.50 | 0.636 |
| Older adults | -0.20 | 0.56 | -0.36 | 0.723 |

Parameter estimates ( $\beta$ ), standard errors (SE), z-values, and false discovery rate (FDR) corrected p-values for 5 ROIs (separated by group) are listed for both young and older adults in each ROI. See Materials and Methods, Fitting model RDMs and brain RDMs for details. Asterisks on the p-value denote an age-related difference. \* $p < 0.05$ . \*\* $p < 0.01$ . \*\*\* $p < 0.001$ .

In young adults, the estimating marginal means revealed that visual representations predicted subsequent image recall in the EVC (95% CI [0.37, 1.64]), LG (95% CI [0.51, 2.02]), FG (95% CI [0.51, 1.85]), and LOC (95% CI [0.45, 1.92]), but not in the vATL (95% CI [−0.24, 1.90]). In older adults, visual representations contributed to subsequent recall memory only in the EVC (95% CI [0.50, 2.46]). No other result was significant (respectively, LG (95% CI [0.03, 2.16]); FG (95% CI [−0.13, 1.72]); LOC (95% CI [−0.93, 1.16]); and vATL (95% CI [−1.09, 1.41])). Although more regions that coded for visual information contributed to later recall memory in young than older adults, post-hoc contrasts between age groups (older adults - young adults) revealed no age-related differences (respectively, β = 0.477, SEM = 0.594, *z* = 0.802, *p* = 0.638 for EVC; β = −0.173, SEM = 0.667, *z* = −0.260, *p* = 0.795 for LG; β = −0.384, SEM = 0.583, *z* = −0.659, *p* = 0.638 for FG; β = −1.070, SEM = 0.652, *z* = −1.641, *p* = 0.504 for LOC; and β = −0.667, SEM = 0.838, *z* = −0.796, *p* = 0.638 for vATL).

In young adults, unlike visual representations, the estimated marginal means revealed that in none of the ROIs semantic representations predicted later recall (respectively, EVC (95% CI [−1.80, 0.40]); LG (95% CI [−0.82, 1.34]); FG (95% CI [−1.33, 0.74]); LOC (95% CI [−1.61, 0.04]); and vATL (95% CI [−1.35, 0.80])). In older adults, in contrast, semantic representations predicted later picture recall in LG (95% CI [1.72, 3.98]) and in FG (95% CI [1.16, 3.30]). No other regions were significant (respectively, EVC (95% CI [−0.60, 1.64]); LOC (95% CI [−1.02, 0.53]); and vATL (95% CI [−1.29, 0.90])). Post-hoc contrasts between age groups (older adults – young adults) revealed that, in LG (β = 2.590, SEM = 0.799, *z* = 3.242, *p* = 0.003) and in FG (β = 2.526, SEM = 0.757, *z* = 3.337, *p* = 0.003), semantic representations predicted subsequent memory to a greater extent in older than in young adults. No other regions showed age-related differences (respectively, β = 1.221, SEM = 0.803, *z* = 1.520, *p* = 0.1214 for EVC; β = 0.543, SEM = 0.575, *z* = 0.945, *p* = 0.431 for LOC; β = 0.073, SEM = 0.782, *z* = 0.094, *p* = 0.925 for vATL).

### Age-related dedifferentiation is compensated by hyperdifferentiation in the fusiform gyrus

To conclude, to examine whether there is an association between regions that showed visual age-related dedifferentiation and regions that showed age-related hyperdifferentation at encoding, supporting successful compensation, we computed an across-subject Pearson’s correlation coefficient between 1) the visual Spearman’s model-brain fit in EVC with the semantic Spearman’s model-brain fit in LG and FG; and 2) the visual Spearman’s model-brain fit in LG with the semantic Spearman’s model-brain fit in LG and FG. In young adults there was a non-significant correlation between EVC and LG, r(16) = 0.11, p = 0.654; and between EVC and FG, r(16) = 0.07, p = 0.769. However, in older adults there was a significant correlation between EVC and FG, r(15) = −0.61, p = 0.009, but not between EVC and LG, r(15) = −0.34, p = 0.186. Moreover, in young adults there was a non-significant correlation between LG and LG, r(16) = 0.41, p = 0.087; and between LG and FG, r(16) = 0.25, p = 0.323. Similarly, in older adults there was a non-significant correlation between LG and LG, r(15) = 0.14, p = 0.009, and between LG and FG, r(15) = −0.17, p = 0.515. The results are shown below in Figure 6.

**Figure 6.**
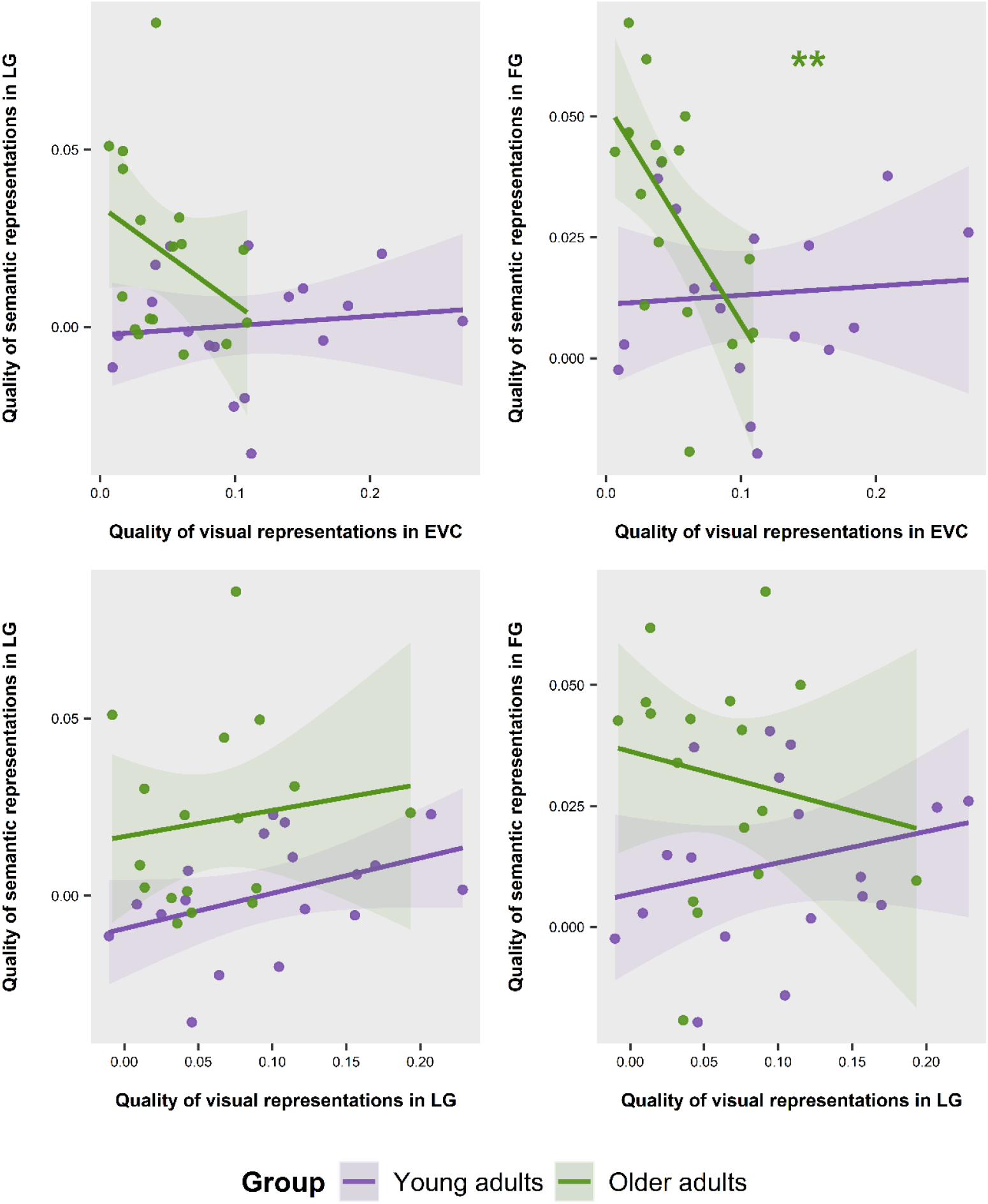
The line plots represent Pearson’s correlation between the visual Spearman’s model-brain fit in EVC and LG with and semantic Spearman’s model-brain in LG and FG for both young (purple line) and older (green line) adults. Purple and green asterisks in the middle of the panels indicate *p* values for significant correlation between ROI. \**p* < 0.05. \*\**p* < 0.01. \*\*\**p* < 0.001.

## Discussion

Our results showed that visual and semantic representations had different strengths in young and older adults and predicted subsequent memory differently in these two groups. By combining models of image properties and prior conceptual knowledge with a task that involved encoding and memory recall, we probed their separate contributions to object encoding. The quality of visual representations was reduced in older adults, consistent with dedifferentiation (Koen and Rugg, 2019), whereas the quality of semantic representations was enhanced in older adults, consistent with hyperdifferentiation (Deng et al., 2021). Turning to subsequent memory, visual representations in the ventral visual pathway predicted later recall of studied objects in young adults. Older adults only showed this effect in the EVC, even though the overall quality of these representations in this group was reduced. In contrast, older adults displayed stronger semantic representations at encoding, which boosted subsequent memory more than in young adults. The data suggest that enhanced semantic processing in aging may boost episodic memory more in older than young adults. Finally, we found that reliance on semantic representations play a compensatory role in the older brain: we observed a negative correlation between visual and semantic representations because older adults with weaker visual representation displayed stronger semantic representations.

Previous studies suggested that aging is associated with neural dedifferentiation (Carp et al., 2011; Koen and Rugg, 2019; Koen et al., 2019, 2020; Trelle et al., 2019), operationalized as reduced specificity of activation patterns for different visual stimuli. Our results revealed that the quality of older adults’ visual representations was reduced compared to young adults in EVC, LG, and vATL. The finding in EVC and LG is in line with the aforementioned study by Deng et al. (2021), which also found impaired visual representations in older adults in EVC. Extending the result of that study, the current study additionally found an age-related reduction in visual representation quality in vATL. Although often considered an amodal semantic hub (Bonner and Price, 2013), vATL is likely to have some degree of modality-specificity (Yi et al., 2007). Indeed, visual objects features were significantly represented in vATL in young adults (see Fig. 4). This finding is consistent with the results of a meta-analysis showing that visual object processing often recruits vATL (Visser et al., 2010). An open question is whether the age-related visual representation deficit we observed in vATL reflects a local effect or it reflects the reduced neural specificity found in EVC, which cascades through the visuo-semantic analyses performed along the ventral pathway, impacting vATL. Even though older adults showed a reduced response towards visual input in EVC and LG at encoding, our findings revealed that they could still use such information at retrieval to recall vivid memories. This finding fits with behavioral results because both groups showed comparable levels of self-report vividness in the scanner, as well as comparable number of hits in the post-scan recognition test. This result suggests that, although neural differentiation compromises the fidelity and efficiency of visual representations (Koen et al., 2020), other factors may play a compensatory role supporting memory in older adults. One of these factors could be an over-reliance on semantic representations.

Our results revealed that semantic representations were stronger in older compared to young adults in the LG and FG. The role of occipito-temporal regions in processing semantic features is well documented in young adults (Koutstaal et al., 2001; Simons et al., 2003; Tyler et al., 2013). Davis et al. (2021) and Naspi et al. (2021) found that some of these regions coded for both visual and semantic information. In particular, emerging data suggest that the FG processes observable (but also verbalizable) semantic features, supporting extraction of meaning from vision. Devereux et al. (2018) combined deep visual and semantic attractor network to model the transformation of vision to semantics, revealing a confluence of late visual and semantic feature representations in FG (see also Tyler et al., 2013). This converges with Martin et al. (2018)’s finding that the FG patterns aligned with the similarity of rated concept words (e.g., hairdryer – comb), a task likely to tap on stored semantic features. Our findings clarify that both image-based visual and semantic features are represented here during successful encoding in both young and older adults. Together, the data further suggest that this initial extraction of visual semantic features is important for the effective encoding of memories of specific objects more in older than young adults.

An important and novel finding of our study is that semantic representations in these occipito-temporal regions also predicted later recall memory in older adults, which is consistent with the results of univariate activation fMRI studies on young adults (Stern et al., 1996; Kirchhoff et al., 2000; Vaidya et al., 2002; Garoff et al., 2005; Kim, 2011). It is unclear why young adults did not benefit from semantic representations as much as the older adults. We believe that age-related differences in representational strength are partly due to differences in strategies used during encoding. Representations are not “hard-wired” into the brain and the representational space can be warped in response to attention (Çukur et al., 2013; Martin et al., 2018; Wang et al., 2018) and therefore encoding strategies. We speculate that the young adults paid more attention to visual features, whereas older adults emphasized more pre-existing semantic features. The latter strategy is also consistent with a greater age-related reliance on gist at encoding, as hypothesized by fuzzy trace theory (Brainerd and Reyna, 2005), and would tend to enhance the fit of activation patterns with the semantic model.

Unlike Deng et al. (2021), the vATL was not engaged in semantic processing by older adults, and consequently it did not contribute to later recall. The lack of engagement of this region may be partly due to stimuli differences; we investigated semantic representations for everyday objects, whereas Deng et al. (2021)’s study examined semantic representations for natural scenes. The semantic properties of a scene (e.g., a farm) are more complex because they contain several objects, which are also likely to appear in other scenes. Thus, to integrate a series of objects into a specific scene, a greater level of semantic abstraction might be required. Thus, it is reasonable to speculate that processing the meaning of scenes is more dependent on the more abstract, interpretive function of vATL than the processing of objects, whose semantic properties could be extracted earlier, in regions such as FG (Bonner and Price, 2013; Lambon Ralph et al., 2017). Further research is required to directly compare the semantic representations of objects and scenes.

Our finding that older adults recruited more anterior ventral pathway regions (e.g., FG) than young adults is consistent with the posterior-anterior shift in aging (PASA; Davis et al., 2008). Evidence for PASA was first reported by Grady et al. (1994) in a positron emission tomography (PET) study that investigated perception of faces and locations. In both conditions, reduced processing efficiency in EVC of older adults led to recruitment of other cortical regions that included, for example, the FG (i.e., area 37) or the prefrontal cortex (PFC; see also Davis et al., 2008). Our results are consistent with these previous finding and more directly suggest that the FG is relevant for semantic processing (Binney et al., 2010; Mion et al., 2010). Although greater semantic processing in older adults may in part explain the absence of age-related differences in recall memory, another factor that is known to account for some of the variance in cognitive ability among older adults is years of education. More years of education have been associated with a slower age-related decline in memory (Colsher and Wallace, 1991; Evans et al., 1993). The results of the cognitive tests suggest that older adults may efficiently use strategies that depend on executive functions in support of their performance in the recall task, especially when the task is cognitively demanding.

In conclusion, visual and semantic representations allow both young and older adults to form memories of specific objects. To our knowledge, this is the first aging fMRI study in which neural representations during encoding are used to predict later recall. Using previously validated representational models, we were able to disentangle visual versus semantic representations. The data supports the idea that aging is associated with neural dedifferentiation of visual features, but it also shows that older adults can rely on stronger semantic representations to boost subsequent memory. This finding challenges the idea that aging is simply associated with a generalized decline in cognitive abilities and their underlying neural mechanisms (e.g., see Fraundorf et al., 2019). Instead, our results revealed that some cognitive and brain mechanisms are not only spared but even enhanced by aging (Umanath and Marsh, 2014). One aspect of cognition that is spared is pre-existing semantic knowledge, but the neural mechanisms of these spared functions were largely unknown. Here, we showed that the LG and FG are part of the semantic network recruited to support memory. We suggested that older adults use this circuit as a compensatory mechanism when the task demand is high like in a recall task.

## Notes

### Competing Interest Statement

The authors have declared no competing interest.

## References

Amunts K, Malikovic A, Mohlberg H, Schormann T, Zilles K (2000) Brodmann’s areas 17 and 18 brought into stereotaxic space - Where and how variable? Neuroimage 11:66–84.

Avants BB, Epstein CL, Grossman M, Gee JC (2008) Symmetric diffeomorphic image registration with cross-correlation: Evaluating automated labeling of elderly and neurodegenerative brain. Med Image Anal 12:26–41.

Beck AT, Ward CH, Mendelson M, Mock J, Erbaugh J (1961) An inventory for measuring depression. Arch Gen Psychiatry 4:561–571 Available at: https://jamanetwork.com/.

Binney RJ, Embleton K v., Jefferies E, Parker GJM, Lambon Ralph MA (2010) The ventral and inferolateral aspects of the anterior temporal lobe are crucial in semantic memory: Evidence from a novel direct comparison of distortion-corrected fMRI, rTMS, and semantic dementia. Cerebral Cortex 20:2728–2738.

Bone MB, Ahmad F, Buchsbaum BR (2020) Feature-specific neural reactivation during episodic memory. Nat Commun 11.

Bonner MF, Price AR (2013) Where is the anterior temporal lobe and what does it do? Journal of Neuroscience 33:4213–4215 Available at: http://www.jneurosci.org/cgi/doi/10.1523/JNEUROSCI.0041-13.2013.

Brainerd CJ, Reyna VF (2005) The science of false memory. Oxford University Press.

Carp J, Park J, Polk TA, Park DC (2011) Age differences in neural distinctiveness revealed by multi-voxel pattern analysis. Neuroimage 56:736–743.

Castel AD (2005) Memory for grocery prices in younger and older adults: The role of schematic support. Psychol Aging 20:718–721.

Castel AD, McGillivray S, Worden KM (2013) Back to the future: Past and future era-based schematic support and associative memory for prices in younger and older adults. Psychol Aging 28:9961003.

Clarke A, Tyler LK (2014) Object-specific semantic coding in human perirhinal cortex. Journal of Neuroscience 34:4766–4775 Available at: http://www.jneurosci.org/content/34/14/4766.short.

Clarke A, Tyler LK (2015) Understanding what we see: How we derive meaning from vision. Trends Cogn Sci 19:677–687 Available at: http://dx.doi.org/10.1016/j.tics.2015.08.008.

Colsher PL, Wallace RB (1991) Longitudinal application of cognitive function measures in a defined population of community-dwelling elders. Ann Epidemiol 1:215–230.

Cox RW, Hyde JS (1997) Software tools for analysis and visualization of fMRI data. NMR Biomed 10:171–178.

Çukur T, Nishimoto S, Huth AG, Gallant JL (2013) Attention during natural vision warps semantic representation across the human brain. Nat Neurosci 16:763–770.

Davis SW, Dennis NA, Daselaar SM, Fleck MS, Cabeza R (2008) Qué PASA? the posterior-anterior shift in aging. Cerebral Cortex 18:1201–1209.

Davis SW, Geib BR, Wing EA, Wang WC, Hovhannisyan M, Monge ZA, Cabeza R (2021) Visual and semantic representations predict subsequent memory in perceptual and conceptual memory tests. Cerebral Cortex 31:974–992.

Deng L, Davis SW, Monge ZA, Wing EA, Geib BR, Raghunandan A, Cabeza R (2021) Age-related dedifferentiation and hyperdifferentiation of perceptual and mnemonic representations. Neurobiol Aging 106:55–67 Available at: https://doi.org/10.1016/j.neurobiolaging.2021.05.021.

Devereux BJ, Clarke A, Tyler LK (2018) Integrated deep visual and semantic attractor neural networks predict fMRI pattern-information along the ventral object processing pathway. Sci Rep 8:1–12 Available at: http://dx.doi.org/10.1038/s41598-018-28865-1.

Du Y, Buchsbaum BR, Grady CL, Alain C (2016) Increased activity in frontal motor cortex compensates impaired speech perception in older adults. Nat Commun 7.

Eickhoff SB, Stephan KE, Mohlberg H, Grefkes C, Fink GR, Amunts K, Zilles K (2005) A new SPM toolbox for combining probabilistic cytoarchitectonic maps and functional imaging data. Neuroimage 25:1325–1335.

Esteban O, Markiewicz CJ, Blair RW, Moodie CA, Isik AI, Erramuzpe A, Kent JD, Goncalves M, DuPre E, Snyder M, Oya H, Ghosh SS, Wright J, Durnez J, Poldrack RA, Gorgolewski KJ (2019) fMRIPrep: a robust preprocessing pipeline for functional MRI. Nat Methods 16:111–116 Available at: http://dx.doi.org/10.1038/s41592-018-0235-4.

Evans DA, Beckett LA, Albert MS, Hebert LE, Scherr PA, Funkenstein HH, Taylor JO (1993) Level of education and change in cognitive function in a community population of older persons. Ann Epidemiol 3:71–77.

Fraundorf SH, Hourihan KL, Peters RA, Benjamin AS (2019) Aging and recognition memory: A meta-analysis. Psychol Bull 145:339–371.

Garoff RJ, Slotnick SD, Schacter DL (2005) The neural origins of specific and general memory: The role of the fusiform cortex. Neuropsychologia 43:847–859.

Grady CL, Maisog JM, Horwitz B, Ungerleider LG, Mentis MJ, Salerno,’ JA, Pietrini P, Wagner E, Haxby’ J v (1994) Age-related Changes in Cortical Blood Flow Activation during Visual Processing of Faces and Location.

Greve DN, Fischl B (2009) Accurate and robust brain image alignment using boundary-based registration. Neuroimage 48:63–72.

Hovhannisyan M, Clarke A, Geib BR, Cicchinelli R, Monge Z, Worth T, Szymanski A, Cabeza R, Davis SW (2021) The visual and semantic features that predict object memory: Concept property norms for 1,000 object images. Mem Cognit 49:712–731 Available at: https://doi.org/10.3758/s13421-020-01130-5.

Jenkinson M, Bannister P, Brady M, Smith S (2002) Improved optimization for the robust and accurate linear registration and motion correction of brain Images. Neuroimage 17:825–841.

Jenkinson M, Smith S (2001) A global optimisation method for robust affine registration of brain images. Med Image Anal 5:143–156 Available at: www.elsevier.com/locate/media.

Kim H (2011) Neural activity that predicts subsequent memory and forgetting: A meta-analysis of 74 fMRI studies. Neuroimage 54:2446–2461 Available at: http://dx.doi.org/10.1016/j.neuroimage.2010.09.045.

Kirchhoff BA, Wagner AD, Maril A, Stern CE (2000) Prefrontal-temporal circuitry for episodic encoding and subsequent memory. Journal of Neuroscience 20:6173–6180.

Koen JD, Hauck N, Rugg MD (2019) The relationship between age, neural differentiation, and memory performance. Journal of Neuroscience 39:149–162.

Koen JD, Rugg MD (2019) Neural dedifferentiation in the aging brain. Trends Cogn Sci 23:547–559.

Koen JD, Srokova S, Rugg MD (2020) Age-related neural dedifferentiation and cognition. Curr Opin Behav Sci 32:7–14.

Koutstaal W, Wagner AD, Rotte M, Maril A, Buckner RL, Schacter DL (2001) Perceptual specificity in visual object priming: Functional magnetic resonance imaging evidence for a laterality difference in fusiform cortex. Neuropsychologia 39:184–199.

Kriegeskorte N, Kievit RA (2013) Representational geometry: Integrating cognition, computation, and the brain. Trends Cogn Sci 17:401–412 Available at: http://dx.doi.org/10.1016/j.tics.2013.06.007.

Kriegeskorte N, Mur M, Bandettini P (2008) Representational similarity analysis – connecting the branches of systems neuroscience. Front Syst Neurosci 2:1–28 Available at: http://journal.frontiersin.org/article/10.3389/neuro.06.004.2008/abstract.

Lambon Ralph MA, Jefferies E, Patterson K, Rogers TT (2017) The neural and computational bases of semantic cognition. Nat Rev Neurosci 18:42–55 Available at: http://dx.doi.org/10.1038/nrn.2016.150.

Martin CB, Douglas D, Newsome RN, Man LLY, Barense MD (2018) Integrative and distinctive coding of visual and conceptual object features in the ventral visual stream. Elife 7:1–29.

McGillivray S, Castel AD (2017) Older and younger adults’ strategic control of metacognitive monitoring: the role of consequences, task experience, and prior knowledge. Exp Aging Res 43:233–256 Available at: http://dx.doi.org/10.1080/0361073X.2017.1298956.

Mion M, Patterson K, Acosta-Cabronero J, Pengas G, Izquierdo-Garcia D, Hong YT, Fryer TD, Williams GB, Hodges JR, Nestor PJ (2010) What the left and right anterior fusiform gyri tell us about semantic memory. Brain 133:3256–3268.

Mishkin M, Ungerleider LG, Macko KA (1983) Object vision and spatial vision: two cortical pathways. Trends Neurosci 6:414–417.

Monge ZA, Madden DJ (2016) Linking cognitive and visual perceptual decline in healthy aging: The information degradation hypothesis. Neurosci Biobehav Rev 69:166–173.

Mumford JA, Turner BO, Ashby FG, Poldrack RA (2012) Deconvolving BOLD activation in event-related designs for multivoxel pattern classification analyses. Neuroimage 59:2636–2643 Available at: http://dx.doi.org/10.1016/j.neuroimage.2011.08.076.

Murray EA, Bussey TJ (1999) Perceptual-mnemonic functions of the perirhinal cortex. Trends Cogn Sci 3:142–151.

Murray EA, Richmond BJ (2001) Role of perirhinal cortex in object perception, memory, and associations. Curr Opin Neurobiol 11:188–193.

Muttenthaler L, Hebart MN (2021) THINGSvision: A python toolbox for streamlining the extraction of activations from deep neural networks. Front Neuroinform 15.

Naspi L, Hoffman P, Devereux B, Morcom AM (2021) Perceptual and semantic representations at encoding contribute to true and false recognition of objects. Journal of Neuroscience 41:8375–8389.

Nasreddine ZS, Phillips NA, Bédirian V, Charbonneau S, Whitehead V, Collin I, Cummings JL, Chertkow H (2005) The Montreal Cognitive Assessment, MoCA: A brief screening tool for mild cognitive impairment. J Am Geriatr Soc 43:695–699 Available at: www.mocatest.

Nili H, Wingfield C, Walther A, Su L, Marslen-wilson W, Kriegeskorte N (2014) A toolbox for representational similarity analysis. PLoS Comput Biol 10:1–11.

Park J, Carp J, Hebrank A, Park DC, Polk TA (2010) Neural specificity predicts fluid processing ability in older adults. Journal of Neuroscience 30:9253–9259.

Penny WD, Trujillo-Barreto NJ, Friston KJ (2005) Bayesian fMRI time series analysis with spatial priors. Neuroimage 24:350–362.

Psychology Software Tools, Inc. [E-Prime 3.0]. (2016).

Reuter M, Rosas HD, Fischl B (2010) Highly accurate inverse consistent registration: A robust approach. Neuroimage 53:1181–1196.

Simons JS, Koutstaal W, Prince S, Wagner AD, Schacter DL (2003) Neural mechanisms of visual object priming: Evidence for perceptual and semantic distinctions in fusiform cortex. Neuroimage 19:613–626.

Simonyan K, Zisserman A (2015) Very deep convolutional networks for large-scale image recognition. In: 3rd International Conference on Learning Representations (Bengio Y, LeCun Y, eds), pp 1–14. San Diego, CA: ICLR 2015. Available at: http://arxiv.org/abs/1409.1556.

Stern CE, Corkin S, González RG, Guimaraes AR, Baker JR, Jennings PJ, Carr CA, Sugiura RM, Vedantham V, Rosen BR (1996) The hippocampal formation participates in novel picture encoding: Evidence from functional magnetic resonance imaging. Proc Natl Acad Sci U S A 93:8660–8665.

Taylor KI, Devereux BJ, Acres K, Randall B, Tyler LK (2012) Contrasting effects of feature-based statistics on the categorisation and basic-level identification of visual objects. Cognition 122:363–374 Available at: http://dx.doi.org/10.1016/j.cognition.2011.11.001.

Trelle AN, Henson RN, Simons JS (2019) Neural evidence for age-related differences in representational quality and strategic retrieval processes. Neurobiol Aging 84:50–60.

Tustison NJ, Avants BB, Cook PA, Zheng Y, Egan A, Yushkevich PA, Gee JC (2010) N4ITK: Improved N3 bias correction. IEEE Trans Med Imaging 29:1310–1320.

Tyler LK, Chiu S, Zhuang J, Randall B, Devereux BJ, Wright P, Clarke A, Taylor KI (2013) Objects and categories: Feature statistics and object processing in the ventral stream. J Cogn Neurosci 25:1723–1735.

Umanath S, Marsh EJ (2014) Understanding how prior knowledge influences memory in older adults. Perspectives on Psychological Science 9:408–426 Available at: http://pps.sagepub.com/lookup/doi/10.1177/1745691614535933.

Vaidya CJ, Zhao M, Desmond JE, Gabrieli JDE (2002) Evidence for cortical encoding specificity in episodic memory: Memory-induced re-activation of picture processing areas. Neuropsychologia 40:2136–2143.

Visser M, Jefferies E, Lambon Ralph MA (2010) Semantic processing in the anterior temporal lobes: A meta-analysis of the functional neuroimaging literature. J Cogn Neurosci 22:1083–1094 Available at: http://www.mitpressjournals.org/doi/10.1162/jocn.2009.21309.

Wang WC, Brashier NM, Wing EA, Marsh EJ, Cabeza R (2018) Neural basis of goal-driven changes in knowledge activation. European Journal of Neuroscience 48:3389–3396.

Yi HA, Moore P, Grossman M (2007) Reversal of the concreteness effect for verbs in patients with semantic dementia. Neuropsychology 21:9–19.

Zeiler MD, Fergus R (2014) Visualizing and understanding convolutional networks. In: European Conference on Computer Vision (Fleet D., ed), pp 813–833. Switzerland: Spring International Publishing.

Zhang Y, Brady M, Smith S (2001) Segmentation of brain MR images through a hidden markov random field model and the expectation-maximization algorithm. IEEE Trans Med Imaging 20:45.

